# Nup358 Sustains Intestinal Epithelial Homeostasis by Preventing Dvl1 Condensate Formation to Restrain Wnt Signaling

**DOI:** 10.64898/2026.03.25.714063

**Authors:** Valeria Guglielmi, Stephen Sakuma, Ethan Y.S. Zhu, Davina Lam, Maximiliano A. D’Angelo

## Abstract

Nucleoporins are increasingly recognized as tissue-specific regulators beyond their structural roles in the nuclear pore complex. Here, we identify nucleoporin Nup358 as a critical repressor of Wnt signaling required for intestinal epithelium integrity. Ablation of Nup358 in adult mice causes a catastrophic loss of crypt-villus architecture and disrupts the intestinal epithelial layer. Notably, while the intestinal stem cell (ISC) pool remains stable, the transit-amplifying (TA) progenitor compartment is depleted. Mechanistically, we show that the interaction of Nup358 with Dvl1 through its N-terminal domain inhibits Dvl1 spontaneous phase separation. In the absence of Nup358, Dvl1 biomolecular condensates promote Tankyrase-mediated degradation of Axin1, leading to the constitutive stabilization of β-catenin and ligand-independent Wnt activation, negatively impacting cell differentiation and TA progenitor survival. Our results demonstrate that Nup358 acts as a molecular safeguard that dampens Wnt signaling levels in intestinal crypts. By preventing Dvl1-mediated Wnt signal amplification, Nup358 allows ISCs to transition into the TA compartment and initiate the differentiation programs essential for intestinal homeostasis.

## Introduction

The intestinal epithelium is a single-cell layer that lines the inner surface of the intestine and plays a key role in maintaining barrier integrity by limiting the entry of luminal antigens and pathogens into the circulation. Because it is constantly exposed to physical and biochemical stress, this epithelium undergoes continuous renewal, with complete turnover occurring every 4-7 days in humans^1^. This rapid regeneration is driven by intestinal stem cells (ISCs) located at the base of intestinal crypts, invaginations of the epithelial layer that form a niche supporting stem cell maintenance and proliferation^2^. ISCs divide asymmetrically to produce either self-renewing stem cells or progenitor cells that enter the transit-amplifying (TA) compartment and begin expansion and lineage commitment. As they move towards the villus, TA progenitors rapidly proliferate and differentiate into specialized epithelial cell types, including enterocytes, goblet cells, and enteroendocrine cells^3^. These cells migrate upward along the villi, where they perform barrier and absorptive functions before being shed into the lumen, completing the renewal cycle. Identifying the factors and mechanisms that regulate intestinal epithelial homeostasis is essential for defining the processes that maintain barrier integrity and for understanding how their dysregulation contributes to disease.

Nuclear pore complexes (NPCs) are large multiprotein channels that span the nuclear envelope connecting the nucleus and the cytoplasm^4^. Besides regulating nucleocytoplasmic transport, NPCs and their components play many transport independent functions^5^. NPCs are composed of 32 different proteins known as nucleoporins^6^. The expression of several nucleoporins varies across cell types, and increasing evidence shows that individual NPC components can perform distinct cell type- and tissue-specific functions^7^. The specialization of nucleoporin function helps explain why alterations in these proteins often result in diseases affecting selective organs^8^. However, the tissue-specific roles of most nucleoporins remain poorly understood, and the mechanisms through which they exert these functions are largely unknown. The nucleoporin Nup358, also known as RanBP2, is a major component of the NPC cytoplasmic filaments^9,10^. Nup358 is a large multidomain protein that contains several Ran-binding domains, a cluster of zinc finger motifs, FG-repeat regions, a cyclophilin homology domain, and a C-terminal SUMO E3 ligase domain^10^. The complex domain organization enables Nup358 to engage in diverse molecular interactions and regulatory activities, and Nup358 has been implicated in a broad range of cellular processes, including nucleocytoplasmic transport, SUMOylation, regulation of mRNA translation, mitotic progression, and microtubule organization among others^11–22^. Additionally, Nup358 dysregulation is linked to various pathologies, including colorectal cancer, blood malignancies, and neurodegenerative disorders^12,23–26^. Despite its critical functions and its connections to disease, the contribution of Nup358 to tissue development and maintenance remains largely unexplored.

Here, we uncover an unexpected role for Nup358 in maintaining intestinal epithelial homeostasis by controlling stem and progenitor cell function. We show that Nup358 ablation in adult mice depletes TA progenitors, leading to the rapid loss of crypt-villus architecture and catastrophic disruption of intestinal epithelium integrity. Mechanistically, Nup358 deficiency promotes the spontaneous formation of Dvl1 condensates, which facilitate Tankyrase-mediated Axin1 degradation. This leads to the constitutive stabilization of β-catenin and aberrant, ligand-independent activation of Wnt signaling, a pivotal pathway regulating intestinal stem cell proliferation and differentiation. Our findings uncover a previously unrecognized nucleoporin-dependent mechanism for restraining canonical Wnt signaling and establish Nup358 as a key modulator of Wnt signaling intensity, safeguarding TA progenitors from Wnt-driven apoptosis and differentiation arrest to maintain epithelial homeostasis.

## RESULTS

### Nup358 ablation disrupts the intestinal epithelium

To determine if Nup358 plays a functional role in tissue development and maintenance *in vivo*, we recently generated a Nup358 mouse knockout line (*Nup358^fl/fl^CreER^T^*^2^, **Extended Data Fig. 1a**)^27^ that allows us to acutely ablate this nucleoporin by tamoxifen treatment (referred to as *Nup358^-/-^* from now on, **Extended Data Fig. 1b**). Notably, acute ablation of Nup358 in adult mice resulted in significant weight loss by 6-7 days post-tamoxifen injection leading to animal mortality (**Fig. 1a**). Analyses of Nup358^-/-^ mice revealed profound abnormalities in the gastrointestinal tract, including enlarged stomach with food retention, and empty small intestines showing significant reduction in both length and luminal cross-sectional area (**Fig. 1b, c**). These findings pointed to alterations in intestinal function and suggested that the weight loss of Nup358^-/-^ mice was caused by defects in nutrient absorption. To investigate if Nup358 ablation affects the integrity of the intestinal tissue, we performed histological analyses of control and Nup358 knockout mice. The intestinal epithelium is organized into two main functional structures: the crypts, which are invaginations at the base of the epithelium housing intestinal stem and progenitor cells, and the villi, which are finger-like projections that extend into the intestinal lumen and are responsible for nutrient uptake (**Extended Data Fig. 1c**). Nup358^-/-^ mice showed extensive alterations in the architecture of the small intestinal epithelium, with disruption of crypt structure and disorganization of the villi epithelial layer (**Fig. 1d**). These findings exposed a role for Nup358 in maintaining the integrity of the intestinal epithelium.

**Figure 1.**
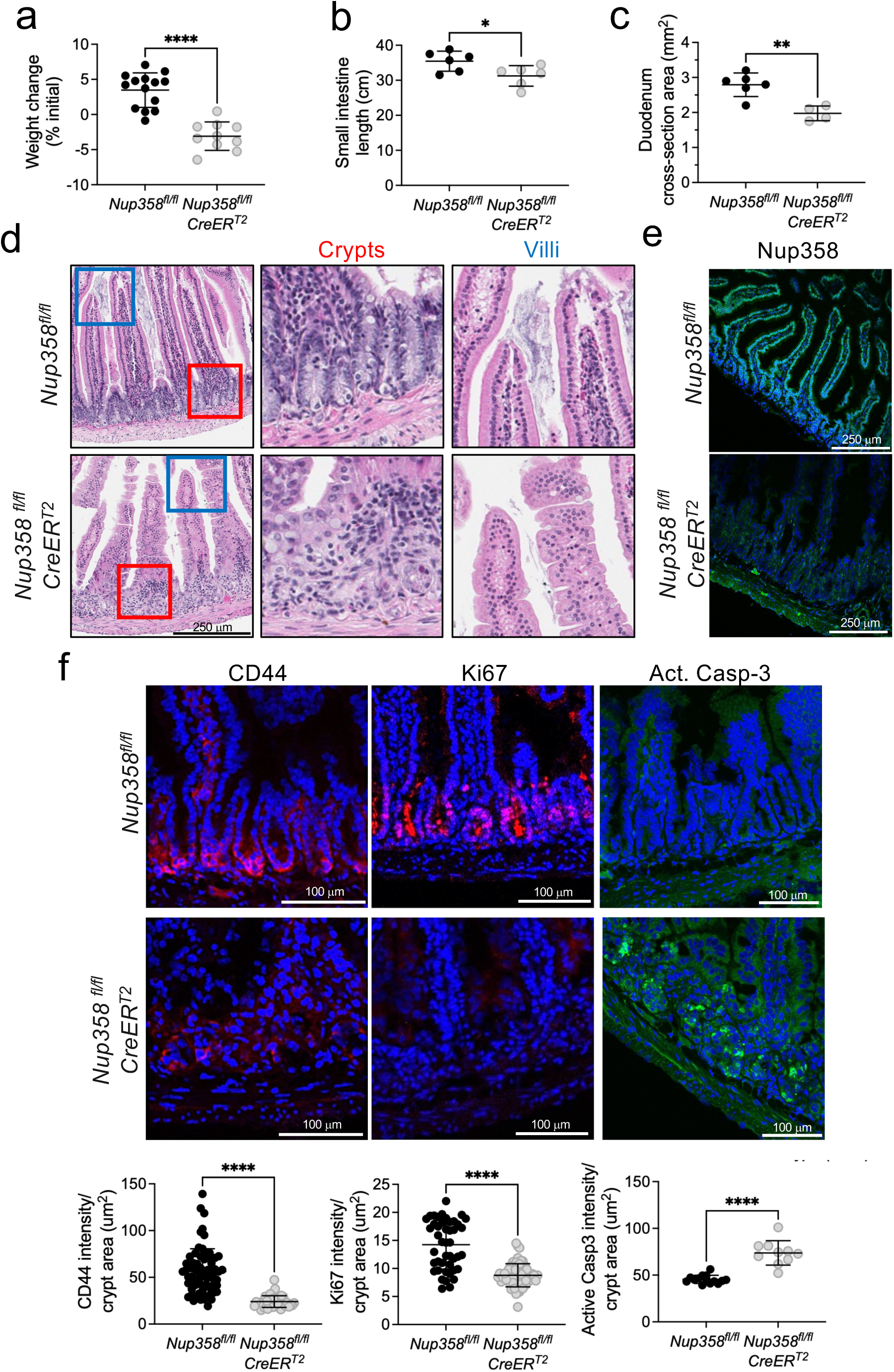
Nup358 ablation disrupts the intestinal epithelium. **a.** Quantification of body weight in *Nup358^fl/fl^*and *Nup358^fl/fl^ CreER^T^*^2^ mice treated with tamoxifen. Data are expressed as mean ± SD. ****p ≤ 0.0001. **b.** Small intestine length in *Nup358^fl/fl^* and *Nup358^fl/fl^ CreER^T2^*mice treated with tamoxifen. Data are expressed as mean ± SD. *p ≤ 0.05. **c.** Duodenum cross-sectional area in *Nup358^fl/fl^* and *Nup358^fl/fl^ CreER^T2^*mice treated with tamoxifen. Data are expressed as mean ± SD. **p ≤ 0.01. **d.** Representative images of hematoxylin and eosin stained small intestine section from *Nup358^fl/fl^*and *Nup358^fl/fl^ CreER^T^*^2^ mice treated with tamoxifen. Color boxes marked the zoom-in regions of crypt and villi. **e.** Immunofluorescence staining for Nup358 in small intestinal epithelium from *Nup358^fl/fl^*and *Nup358^fl/fl^ CreER^T^*^2^ mice treated with tamoxifen. **f.** Immunofluorescence staining for CD44, Ki67, and activated caspase 3 in small intestinal epithelium from *Nup358^fl/fl^* and *Nup358^fl/fl^ CreER^T^*^2^ mice treated with tamoxifen. Bottom: Quantification of immunofluorescence intensity per area for CD44, Ki67, and activated caspase 3.

### Nup358 knockout does not affect the number of ISCs but strongly reduces TA progenitor cells

The crypts are mainly occupied by undifferentiated cells, including ISCs and TA progenitors, that continuously divide to replenish the lining of the intestine. In homeostatic conditions, ISCs produce TA progenitor cells, which rapidly proliferate as they move towards the villus (**Extended Data Fig. 1c**). As TA cells move away from the crypt and into the villus, they begin to differentiate into the different types of intestinal epithelial cells, including absorptive enterocytes, goblet cells, and enteroendocrine cells, that migrate upward to maintain the epithelial barrier and digestive function. Immunofluorescence analyses of small intestine sections showed that Nup358 is expressed similarly in crypt and villus cells (**Fig. 1e**). To investigate if Nup358 ablation affects crypt structure by disrupting the homeostasis of the intestinal stem and progenitor cells, we stained small intestine sections with cell-type-specific markers. We found that immunostaining for CD44, which labels both ISCs and TA progenitors, was strongly reduced in the crypts of Nup358 knockout animals compared to controls (**Fig. 1f, g**). The intestinal epithelium of Nup358^-/-^ mice also displayed reduced expression of the proliferation marker Ki67 and higher levels of the apoptosis marker activated caspase 3 (**Fig. 1f, g, Extended Data Fig. 1d**). TA cells are the most proliferative compartment of the intestinal crypt, and Ki67 is generally used as a marker for this population. Therefore, the reduced levels of Ki67 staining support the loss of TA progenitors. Because CD44 does not distinguish between ISCs and TA progenitors, we stained small intestine sections against Olfm4, an ISC-specific marker. A similar intensity of Olfm4 staining was detected between control and Nup358^-/-^ mice, indicating that the number of ISCs was not affected by Nup358 ablation (**Fig. 2a**). However, while in control mice Olfm4⁺ ISCs were restricted to the crypt base, in Nup358^-/-^ mice these cells were dispersed throughout the crypt region, reflecting a loss of intestinal tissue organization. These findings indicate that Nup358 is dispensable for the maintenance of ISCs but plays a critical role in sustaining the downstream TA progenitors.

**Figure 2.**
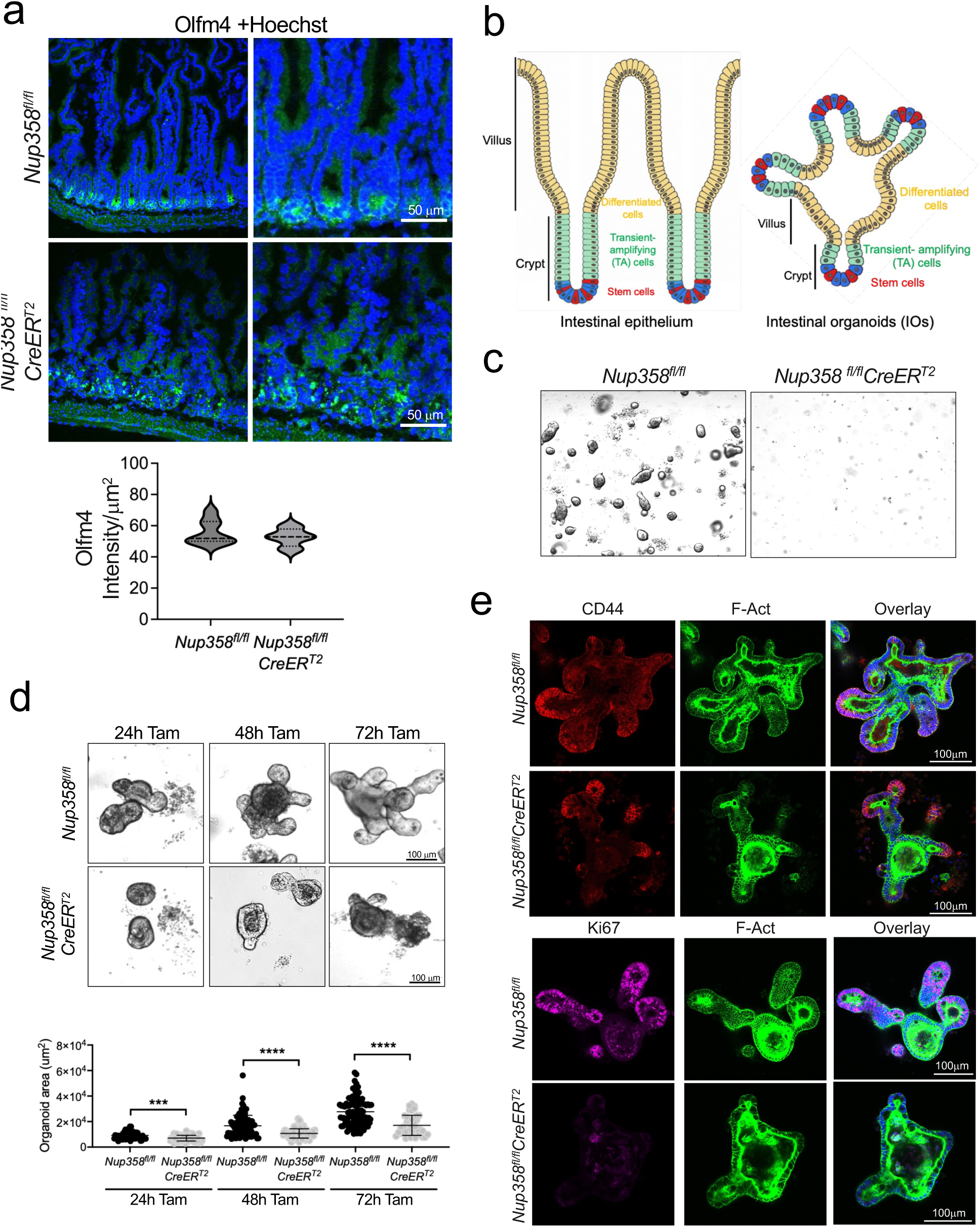
Nup358 knockout results in the loss of transient-amplifying (TA) progenitors without affecting intestinal stem cells (ISCs). **a.** Immunofluorescence staining for Olfm4 in small intestinal epithelium from *Nup358^fl/fl^* and *Nup358^fl/fl^ CreER^T^*^2^ mice treated with tamoxifen. Bottom: Quantification of Olfm4 immunofluorescence intensity per area. **b.** Schematic illustration of the structural organization of the small intestinal epithelium (left) and intestinal organoids (right). **c.** Intestinal organoids (IOs) obtained from *Nup358^fl/fl^* and *Nup358^fl/fl^ CreER^T^*^2^ mice treated with tamoxifen. **d.** Crypts isolated from *Nup358^fl/fl^* and *Nup358^fl/fl^ CreER^T^*^2^ mice were cultured in IO conditions for 24 hours before treating them with tamoxifen for 24, 48, and 72 hours. Bottom: Quantification of intestinal organoid area. Data are expressed as mean ± SD, ***p ≤ 0.001, ****p ≤ 0.0001. **e.** Immunofluorescence staining for CD44 and Ki67 in IOs from *Nup358^fl/fl^* and *Nup358^fl/fl^ CreER^T^*^2^ mice cultured for 7-9 days, before tamoxifen treatment with tamoxifen for 24 hours (CD44) or 72 hours (Ki67). IOs were counterstained with phalloidin.

### Nup358 is required for ISc/progenitor function

Despite having normal numbers of ISCs, Nup358 knockout mice exhibit a marked reduction in CD44⁺ and Ki67⁺ cells and increased activated caspase 3 staining (**Fig. 1f, g**). These findings suggest that Nup358 is required for the differentiation of ISCs into TA progenitors and/or for TA cell survival. Intestinal organoids (IOs) closely recapitulate the structure, cell diversity, and regenerative dynamics of the intestinal epithelium and are a well-established model to study ISC function^28,29^. IOs can be generated from isolated crypts or individual ISCs, that self-organize into crypt- and villus-like domains, mimicking the architecture of the intestinal epithelium (**Fig. 2b**). To investigate the effect of Nup358 ablation on ISC function, we established IOs from crypts isolated from intestines of tamoxifen treated control *Nup358^fl/fl^* and *Nup358^fl/fl^CreER^T^*^2^ mice. Ablation of Nup358 prior to crypt isolation completely abolished organoid formation, confirming a requirement for Nup358 for ISC function (**Fig. 2c**). To circumvent this early block and assess whether Nup358 depletion affects the growth and differentiation of established organoids, we isolated intestinal crypts from untreated *Nup358^fl/fl^CreER^T^*^2^ *and Nup358^fl/fl^* control mice and cultured them under IO conditions for 24 hours before tamoxifen treatment. Twenty-four hours after tamoxifen treatment (48 hours of organoid formation), organoids already showed delayed growth and impaired crypt and villi development, which was even more pronounced at 48 and 72 hours post-tamoxifen treatment **(Fig. 2d, Extended Fig. 1e**). Live imaging of organoids starting at 96 hours post-tamoxifen showed markedly reduced growth, crypt development, and villi formation in Nup358^-/-^ organoids compared to controls and resulted in premature organoid degeneration **(Extended Data Videos 1, 2**). Moreover, when Nup358 was ablated after 7-9 days of culture, when organoids had developed well-defined crypt and villus structures, a striking reduction in CD44 and Ki67 staining was observed (**Fig. 2e**), consistent with our *in vivo* observations. Together, these findings support a critical role for Nup358 in ISC function and the maintenance of TA progenitors.

### Nup358 ablation activates Wnt signaling

Wnt signaling is a major regulator of intestinal epithelial homeostasis with a critical role in ISC self-renewal and proliferation^30–33^. Notably, we found an upregulation of the canonical Wnt targets *Axin2*, *Ccnd1*, and *cMyc*, in the small intestinal epithelium of Nup358^⁻/⁻^ mice (**Fig. 3a**), suggesting activation of this pathway. Under physiological conditions, Wnt activity is highest at the crypt base and gradually declines toward the villus tip, forming a gradient that is crucial for maintaining intestinal epithelial homeostasis **(Extended Data Fig. 1c**). As ISC daughters enter the TA compartment, the progressive decline in Wnt signaling enables their exit from the proliferative stem cell state and initiates differentiation. Disruption of this gradient or inability to shut down Wnt signaling can impair TA cell expansion by blocking ISC differentiation and triggering apoptosis^33–36^. To further confirm that Wnt signaling is altered by Nup358 ablation, we analyzed the levels and localization of the β-catenin partner TCF-4 in intestinal sections of control and Nup358^-/-^ mice. TCF-4 showed higher levels and increased nuclear accumulation in Nup358^-/-^ mice, particularly in the more differentiated regions of the intestinal epithelium towards the villus domain (**Fig. 3b, Extended Data Fig. 2a**). These findings show that Wnt signaling is dysregulated in the absence of Nup358. To investigate how Nup358 modulates Wnt activity, we generated a stable HEK293T cell line expressing the 7TGP fluorescent Wnt reporter, in which GFP expression is driven by seven TCF/LEF regulatory elements^37^. This reporter allows sensitive detection of canonical β-catenin-dependent pathway activation. HEK293T cells expressing 7TGP were transfected with control or Nup358-targeting siRNAs, and GFP expression was imaged by confocal microscopy and quantified by flow cytometry. Notably, Nup358 downregulation led to a robust increase in GFP levels in the absence of Wnt stimulation, indicating aberrant ligand-independent activation of the Wnt pathway (**Fig. 3c**). Similar activation of the reporter was observed when Nup358 was downregulated with shRNAs (**Extended Data Fig. 2b, c**). To confirm that this activation was β-catenin-dependent, we depleted β-catenin in Nup358-deficient cells using specific siRNAs. β-catenin knockdown strongly suppressed Nup358 knockdown-mediated Wnt reporter activation (**Fig. 3d**). Together, these data demonstrate that Nup358 downregulation induces β-catenin-dependent Wnt signaling, indicating that this nucleoporin functions as a negative regulator of the pathway.

**Figure 3.**
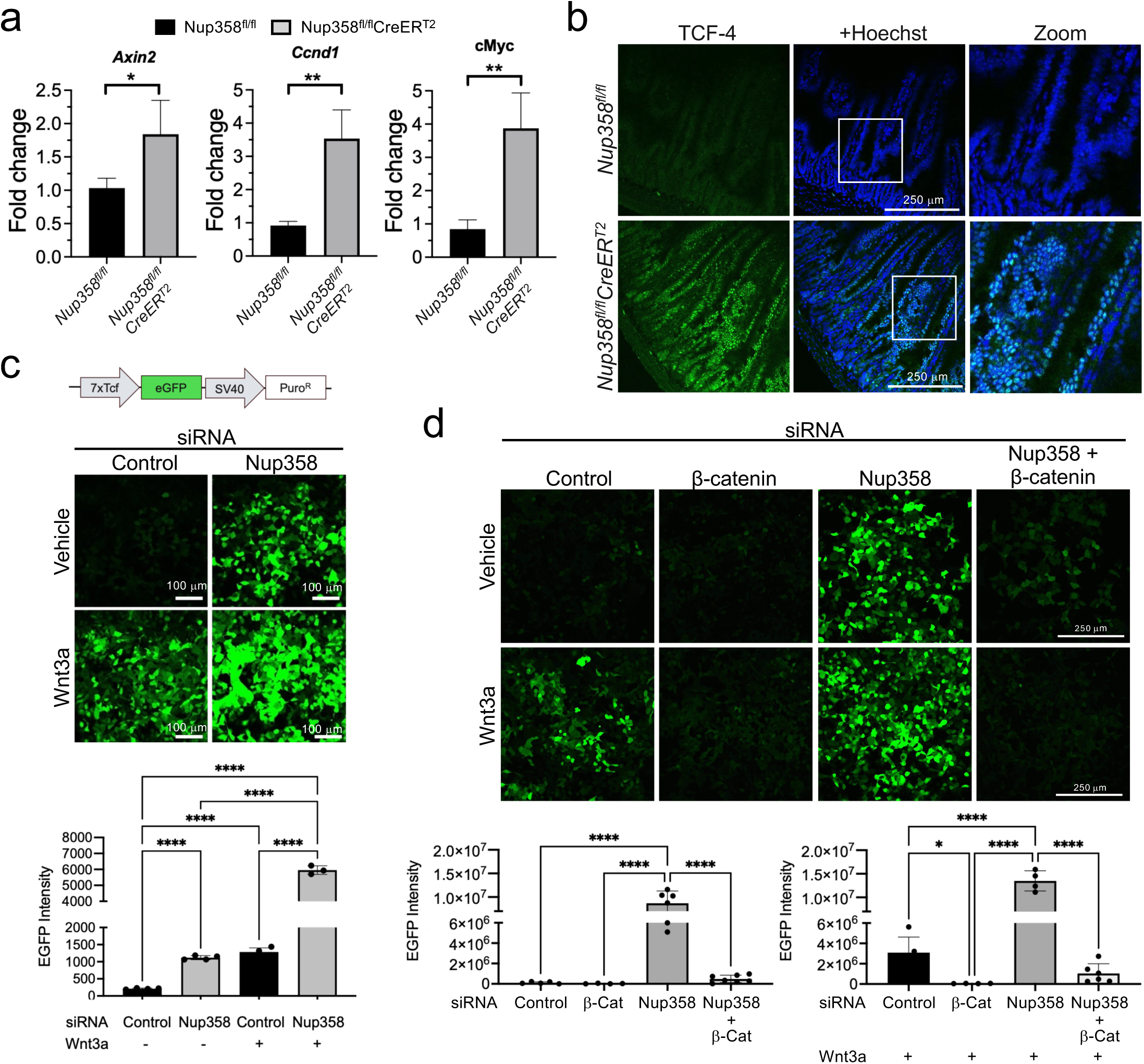
Nup358 represses Wnt signaling pathway. **a.** The expression levels of Wnt target genes Axin2, Ccnd1, and cMyc were analyzed in small intestinal epithelium from *Nup358^fl/fl^*and *Nup358^fl/fl^ CreER^T^*^2^ mice treated with tamoxifen by qPCR. Data are expressed as mean ± SD. *p ≤ 0.05 and **p ≤ 0.01. **b.** Immunofluorescence staining for TCF-4 in small intestinal epithelium from Nup358^fl/fl^ and *Nup358^fl/fl^ CreER^T^*^2^ mice treated with tamoxifen. **c.** Fluorescent Wnt reporter activity in HEK293T cells transfected with either control or Nup358-specific siRNA and treated with either Wnt3a or vehicle were analyzed by confocal microscopy (top) and quantified by flow cytometry (bottom). Data are expressed as mean ± SD, ****p ≤ 0.0001. **d.** Fluorescent Wnt reporter activity in HEK293T cells transfected with control siRNA, β-catenin-specific siRNA, Nup358-specific siRNA, or a combination of Nup358-specific siRNA and β-catenin-specific siRNA and treated with either Wnt3a or vehicle were analyzed by confocal microscopy (top) and quantified (bottom). Data are expressed as mean ± SD. *p ≤ 0.05 and ****p ≤ 0.0001.

### Nup358 inhibition leads to the formation of cytoplasmic Dvl1 condensates

Canonical Wnt signaling functions primarily through the stabilization of β-catenin^38^. In resting conditions, β-catenin is kept at low levels through continuous degradation by the destruction complex, which includes Axin, APC, GSK3β, and CK1α. This complex phosphorylates β-catenin leading to its ubiquitination and proteasomal degradation^39^. Binding of Wnt ligands to Frizzled (Fzd) receptors and LRP5/6 co-receptors recruits and activates the cytoplasmic Dvl proteins, which recruit Axin to the plasma membrane, disassembling the destruction complex and preventing β-catenin degradation^40^. The accumulation of β-catenin and translocation into the nucleus activates Wnt target genes through TCF/LEF transcription factors. Our data shows that Nup358 downregulation is sufficient to activate Wnt signaling in a ligand-independent manner (**Fig. 3c**). We also found that Nup358-depleted cells display stronger activation when treated with the Wnt3a ligand (**Fig. 3c, d**). This additive effect implies that ligand-mediated activation is still functional in the absence of Nup358 and suggests that this nucleoporin represses Wnt downstream of the receptor complex. In vertebrates, there are three Dvl paralogs, Dvl1, Dvl2, and Dvl3^41^. A previous study identified an interaction between Nup358 and Dvl1 and suggested that Nup358 modulates neuronal polarization by negatively regulating Dvl1 activity^42^. To determine if Nup358-mediated repression of Wnt signaling involves Dvl1, we examined if its downregulation is sufficient to block Wnt activation in Nup358-deficient cells. For this we downregulated each protein individually or in combination and analyzed the activity of the 7TGP Wnt reporter in HEK293T cells. Given the essential role of Dvl1 in ligand-mediated Wnt pathway activation, Dvl1 knockdown strongly suppressed 7TGP reporter activation in response to Wnt3a stimulation, but it had no effect in unstimulated conditions (**Fig. 4a, Extended Data Fig. 3a, b**). Notably, co-depletion of Dvl1 and Nup358 markedly impaired 7TGP Wnt reporter activation, indicating that Dvl1 mediates the Wnt signaling activation observed in Nup358-deficient cells (**Fig. 4a, Extended Data Fig. 3b**).

**Figure 4.**
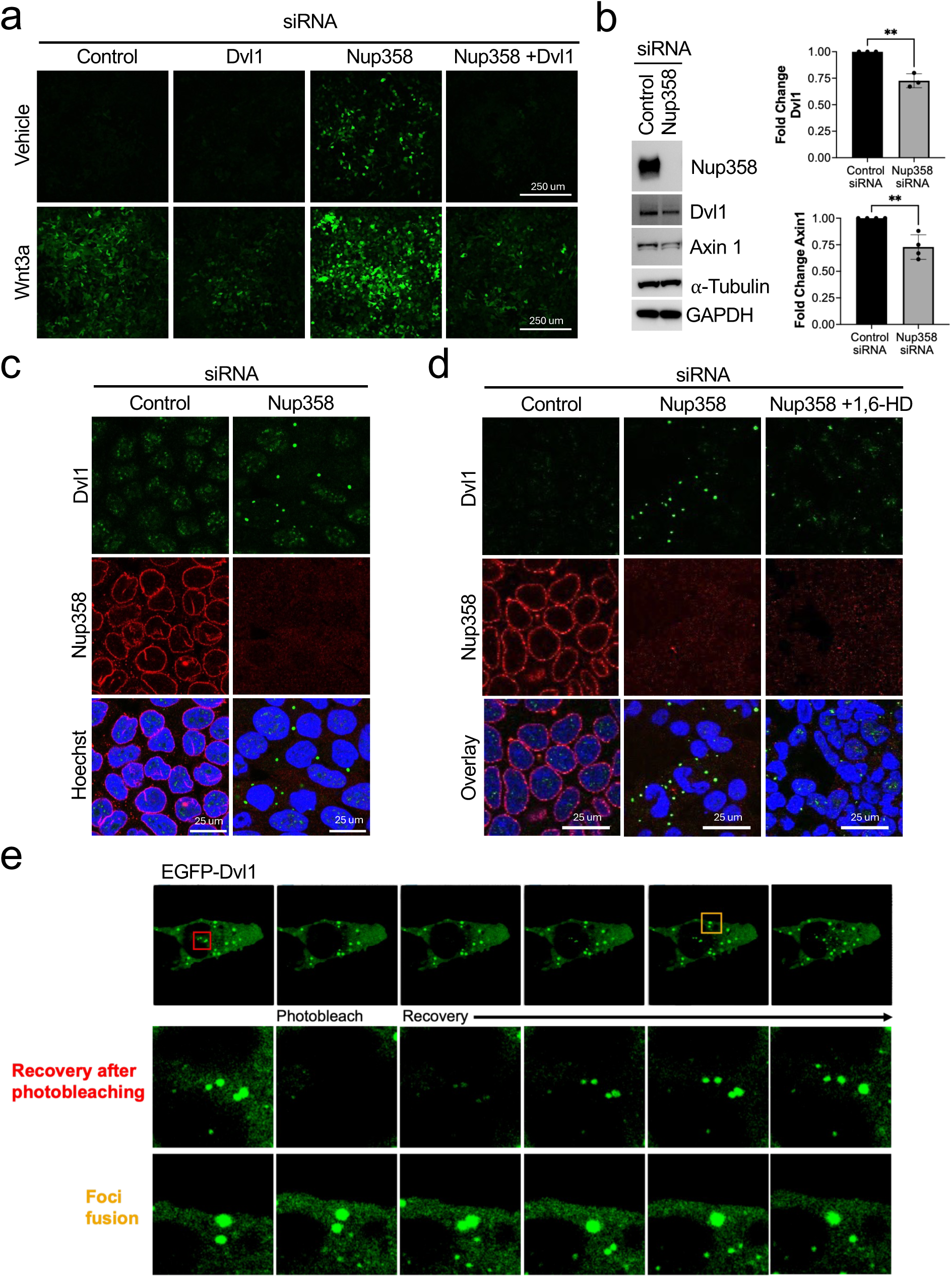
Nup358 inhibition activates Wnt pathway by promoting the formation of cytoplasmic Dvl1 condensates. **a.** Levels of the 7TGP fluorescent Wnt reporter in HEK293T cells transfected with control siRNA, Dvl1-specific siRNA, Nup358-specific siRNA, or a combination of Nup358-specific siRNA and Dvl1-specific siRNA and treated with either Wnt3a or vehicle. **b.** Western blot analysis and relative quantification of protein levels of Dvl1 and Axin1 in HEK293T cells transfected with either control or Nup358-specific siRNA. α-Tubulin and GAPDH were used as loading controls. Data are expressed as mean ± SD. **p ≤ 0.01. **c.** Immunofluorescence staining for Dvl1 and Nup358 in HEK293T cells transfected with either control or Nup358-specific siRNA. **d.** Immunofluorescence staining for Dvl1 and Nup358 in HEK293T cells transfected with control or Nup358-specific siRNA and treated with vehicle or 1,6-hexanediol 5% for 2 minutes. **e.** Fluorescence recovery after photobleaching (FRAP) and fusion analysis of Dvl1 condensates in HEK293T cells transfected with Nup358-specific siRNA and EGFP-Dvl1.

Wnt/β-catenin signaling is regulated by the dynamic assembly of multiple biomolecular condensates^43–48^. Upon engagement of Fzd and LRP5/6 receptors by Wnt ligands, Dvl proteins multimerize and undergo liquid-liquid phase separation (LLPS), concentrating key pathway components such as Axin and LRP6 into localized clusters ^46,49^. These Dvl condensates, known as signalosomes, promote receptor clustering, recruit downstream effectors, and disrupt the formation of the β-catenin destruction complex by sequestering Axin. This prevents β-catenin phosphorylation and degradation, promoting the activation of Wnt target genes. Notably, while Dvl1 total protein levels were slightly reduced by Nup358 downregulation, analysis of its intracellular localization showed the formation of large Dvl1 foci (**Fig. 4b, c**). These foci localized to the cytoplasm rather than the plasma membrane, indicating that Wnt activation in Nup358-depleted cells is not driven by increased receptor concentration. This is consistent with the ligand-independent Wnt activation observed in these cells (**Fig. 3c, d**). Similar Dvl1 foci formed in HT-29 colorectal cancer cells depleted of Nup358, which also showed increased nuclear accumulation of TCF-4 (**Extended Data Fig. 3c, d**). To determine if the Dvl1 aggregates represent condensates, cells transfected with control or Nup358-specific siRNAs were incubated with 1,6-hexanediol (1,6-HD), an alcohol that disrupts liquid-liquid phase separation and dissolves biomolecular condensates, before fixation and immunofluorescence. Treatment with 1,6-HD strongly decreased foci formation indicating that Dvl1 foci in Nup358-depleted cells represent biomolecular condensates (**Fig. 4d**). To further confirm this, Nup358 was downregulated in 293T cells expressing EGFP-Dvl1 and foci formation was analyzed by live fluorescent microscopy. In these cells, EGFP-Dvl1 formed highly dynamic, condensate-like foci that exhibited frequent fusion events and showed rapid fluorescence recovery after photobleaching (**Fig. 4e, Extended Data Video 2**). Our findings indicate that Nup358 restrains the self-association of Dvl1, blocking its assembly into cytoplasmic condensates.

### Nup358 represses Wnt signaling by preventing Tankyrase-mediated Axin1 degradation in Dvl1 condensates

Canonical Wnt signaling is controlled by degradation of β-Catenin. Axin1 functions as the core scaffold of the β-catenin degradation complex that brings together all the key components^40,50^. In unstimulated conditions, Axin1 is protected from Tankyrase-mediated degradation through phosphorylation by CK1α, which is concentrated by the formation of degradation complex condensates. The sequestration of Axin1 by Dvl1 during signalosome formation allows Tankyrase to modify Axin1 promoting its degradation and disrupting the formation of the β-catenin destruction complex^51^. Since Nup358 downregulation leads to large cytoplasmic Dvl1 condensates, we speculated that the aberrant Dvl1 aggregation could lead to Axin1 degradation and β-catenin stabilization in a ligand-independent manner. Supporting this hypothesis, Nup358-depleted cells show lower Axin1 levels and higher β-catenin levels (**Fig. 4b, Extended Data Fig. 4a**). To investigate if Nup358 downregulation inhibits β-catenin protein degradation, we ectopically expressed β-catenin-GFP in HEK293T cells under the control of a CMV promoter and subsequently transfected them with control or Nup358-specific siRNAs. This approach allowed us to determine the effect of Nup358 downregulation on β-catenin protein levels independently of its endogenous transcriptional regulation. In controls, a very small number of cells showed GFP signal, consistent with β-catenin protein being continuously degraded (**Fig. 5a**). Contrarily, Nup358-depleted cells showed a strong increase in β-catenin-GFP positive cells, which also showed higher GFP intensity, confirming the inhibition of its degradation pathway (**Fig. 5a**).

**Figure 5.**
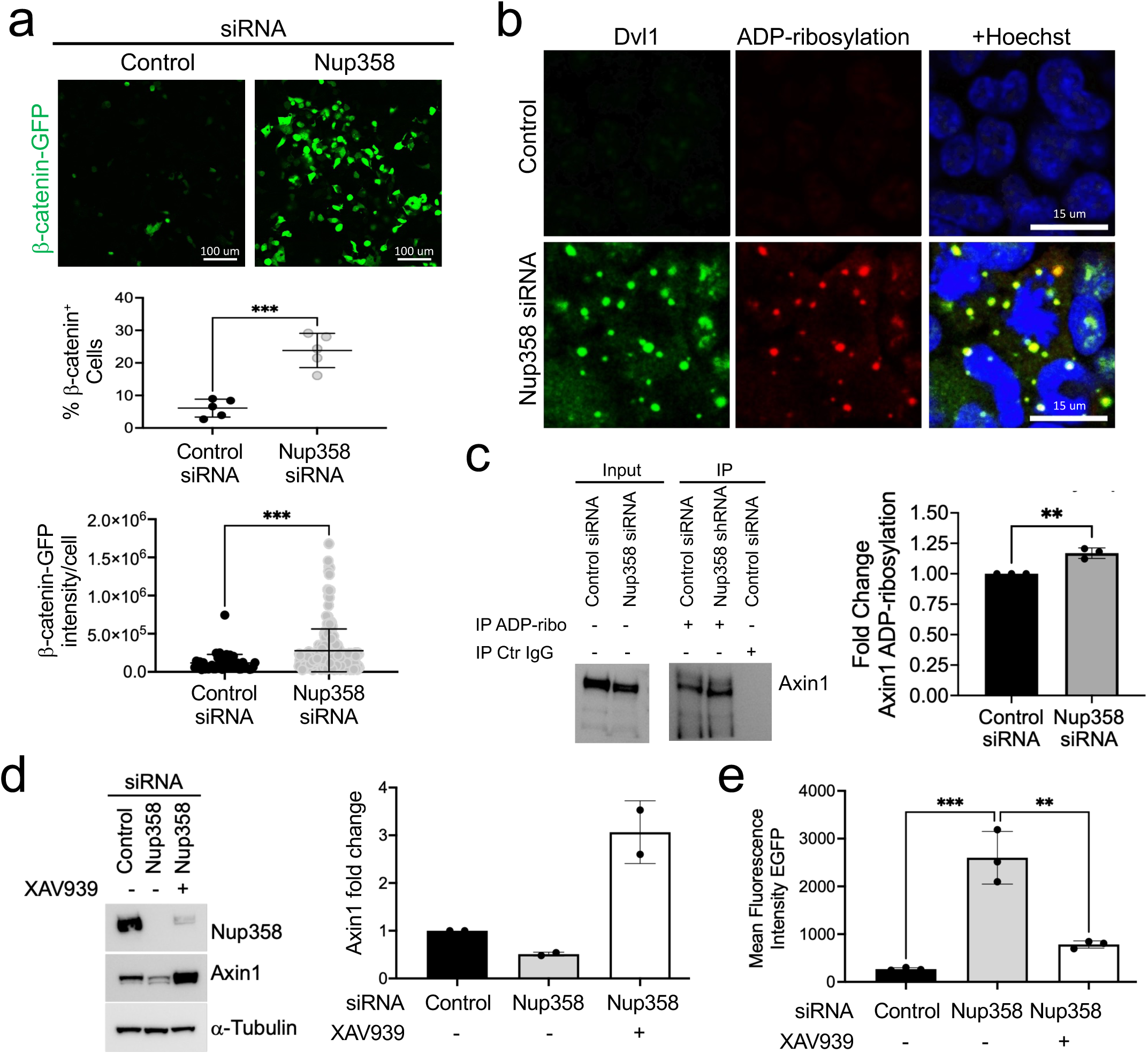
Nup358 depletion inhibits β-catenin degradation by inducing Tankyrase-mediated degradation of Axin1. **a.** HEK293T cells were transfected with β-catenin-GFP plasmid and control or Nup358-specific siRNAs and GFP expression was analyzed by confocal microscopy. The percentage of GFP-positive cells and GFP intensity per cell were quantified. Data are expressed as mean ± SD. ***p ≤ 0.001. **b.** Immunofluorescence staining for Dvl1 and ADP-ribosylation in HEK293T cells transfected with either control or Nup358-specific siRNAs. **c.** Western blot analysis of co-immunoprecipitation of ADP-ribosylation and Axin1 in HEK293T cells transfected with control or Nup358-specific siRNA. Axin1 levels in immunoprecipitates were quantified relative to controls and normalized to Axin1 input levels. Data are expressed as mean ± SD. **p ≤ 0.01 **d.** Left: Western blot analyses of Nup358 and Axin1 in HEK293T cells transfected with control or Nup358-specific siRNA and treated with 200 nM XAV-939 or vehicle for 72 hours. α-Tubulin was used as loading control. Right: Quantification of Axin1 protein levels (n=2 experiments). **e.** Quantification of fluorescent Wnt reporter activity by flow cytometry analysis in HEK293T cells transfected with control siRNA or Nup358-specific siRNA and treated with 200nM XAV-939 or vehicle for 72 hours. Data are expressed as mean ± SD. **p ≤ 0.01 and ***p ≤ 0.001.

Axin1 degradation is promoted by Tankyrase-mediated ADP-ribosylation, which facilitates its ubiquitination and proteasomal degradation^51^. Notably, ADP-ribose staining revealed strong signal in cytoplasmic foci that colocalize with Dvl1 puncta in Nup358-depleted cells (**Fig. 5b, Extended Data Fig. 4b**). Moreover, immunoprecipitation with an anti-ADP-ribosylation antibody pulled down higher levels of Axin1, indicating an increase in its ADP-ribosylation (**Fig. 5c**). These findings suggest that the abnormal aggregation of Dvl1 may promote Axin1 degradation through Tankyrase, leading to stabilization of β-catenin. If this is the case, we could expect that Tankyrase inhibition should restore Axin1 levels and prevent Wnt activation in Nup358 knockdown cells. To test this, HEK293T cells expressing the 7TGP Wnt reporter were transfected with control and Nup358-specific siRNAs and subsequently treated with vehicle or the Tankyrase-specific inhibitor XAV939^51^. We found that Tankyrase inhibition restored Axin1 levels in Nup358-depleted cells (**Fig. 5d**) and strongly suppressed Wnt pathway activation (**Fig. 5e**). Tankyrase inhibition resulted in a partial reduction of the global ADP-ribosylation signal, consistent with the fact that not all cellular ADP-ribosylation events are Tankyrase-dependent (**Extended Data Fig. 4c**). Also, the decrease in ADP-ribosylation did not affect the formation of Dvl1 cytoplasmic foci, indicating that this modification is not required for Dvl1 aggregation (**Extended Data Fig. 4d**). Together, these data suggest that Nup358 stabilizes Axin1 by limiting Tankyrase-dependent modification and degradation, thereby restraining Wnt signaling.

### Nup358 inhibits Dvl1 condensate formation through its N-terminal leucine-rich domain

Nup358 is a multifunctional nucleoporin that contains several distinct functional domains, including a leucine-rich domain, Ran binding domains, phenylalanine-glycine (FG) repeats, internal disordered domains, and a SUMO E3 ligase domain and a cyclophilin-like domain (**Fig. 6a**)^10^. Nup358 works together with Ubc9 and RanGAP1-SUMO1 to SUMOylate many cellular targets and many of its functions are modulated though its E3 SUMO ligase activity^19,52^. Nup358-downregulation leads to loss of Ubc9 from nuclear pores (**Extended Data Fig. 5a**)^21^, and Ubc9 ablation in mice was found to also affect intestinal homeostasis^53^. Moreover, previous studies linked Nup358-dependent SUMOylation of APC and TCF-4 to regulation of β-catenin activity, although this modification was found to have a positive effect on Wnt signaling^54,55^. To determine whether the loss of Nup358-mediated SUMOylation is responsible for Wnt activation, we used two complementary approaches. First, we inhibited SUMOylation in HEK293T cells expressing the 7TGP Wnt reporter by downregulating Ubc9, the only cellular SUMO E2 ligase^56^, using specific siRNAs that we previously validated (**Extended Data Fig. 5b**)^21^. Despite the efficient loss of Ubc9, this treatment failed to activate Wnt signaling (**Fig. 6b**). Second, we inhibited SUMOylation with the selective inhibitor 2-D08, which blocks the transfer of SUMO peptide to substrates^57^. This treatment also did not activate the 7TGP Wnt reporter (**Fig. 6b**). These findings indicate that Nup358 does not repress Wnt activity through SUMOylation.

**Figure 6.**
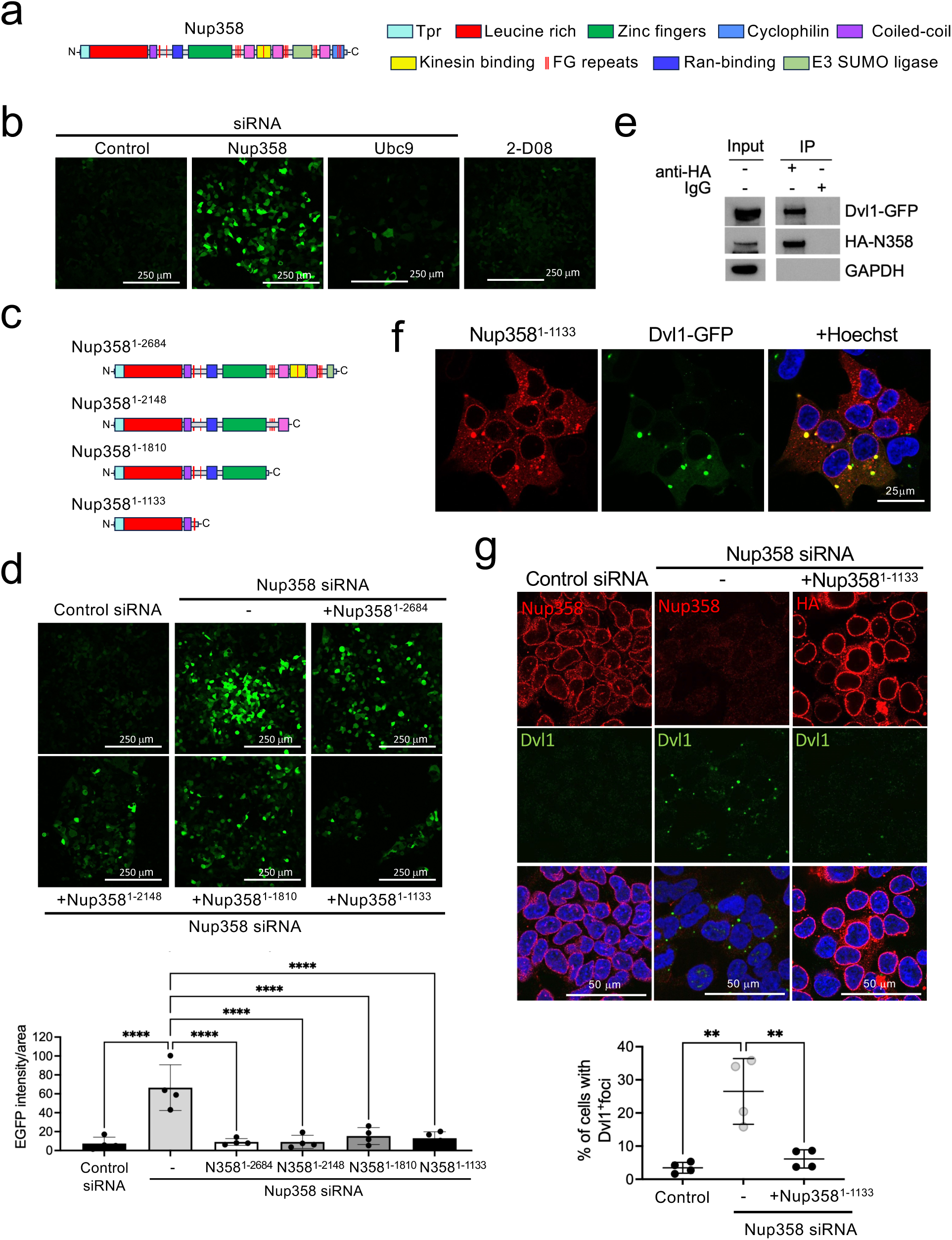
Nup358 inhibits Dvl1 condensate formation through its N-terminal leucine-rich domain. **a.** Schematic representing different functional domains of Nup358. **b.** Fluorescent Wnt reporter activity in HEK293T cells transfected with control siRNA, Nup358-specific siRNA, Ubc9-specific siRNA, or treated with the SUMOylation inhibitor 2-D08. **c.** Schematic representing different HA-tagged truncated forms of Nup358 that were transiently expressed in HEK293T cells. **d.** Fluorescent Wnt reporter activity in HEK293T cells transfected with either control or Nup358-specific siRNA alone or in combination with different HA-tagged truncated forms of Nup358 were analyzed by confocal microscopy (top) and quantified (bottom). Data are expressed as mean ± SD. ****p ≤ 0.0001 **e.** Western blot analysis of co-immunoprecipitation of HA-Nup358 and EGFP-Dvl1 transiently expressed in HEK293T cells. GAPDH was used as loading control. **f.** Immunofluorescence of HEK293T cells transfected with HA-Tagged Nup358^1–1133^ and EGFP-Dvl1and stained with an anti-HA antibody. **g.** Immunofluorescence staining for endogenous Dvl1 and Nup358 in HEK293T cells transfected with either scramble control or Nup358-specific siRNA alone or in combination with HA-Tagged Nup358^1–1133^ fragment were analyzed by confocal microscopy (top) and quantified (bottom). Data are expressed as mean ± SD. **p ≤ 0.01.

To identify the Nup358 domain responsible for repressing Dvl1 condensate formation, we expressed HA-tagged truncation mutants of this nucleoporin (**Fig. 6c, Extended Data Fig. 5c**) and assessed whether specific domains could suppress the aberrant activation of the 7TGP Wnt reporter in Nup358-downregulated cells. (**Fig. 6d**). Consistent with SUMOylation being dispensable for the repression of Wnt signaling, mutants lacking the E3 SUMO ligase domain were still able to repress 7TGP activation (**Fig. 6d**). The same was observed with deletion of the Ran binding, Zinc finger, and cyclophilin homology domains, and expression of the Nup358 N-terminal leucine-rich region was sufficient to rescue Wnt repression in Nup358-depleted cells (**Fig. 6d**). The N-terminal domain of Nup358 was previously found to associate with Dvl1 in neurons. Consistent with these findings, immunoprecipitation experiments revealed that the Nup358 N-terminal domain interacts with Dvl1 (**Fig. 6e**) and also localizes to biomolecular condensates induced by ectopic expression of Dvl1-GFP (**Fig. 6f**). Moreover, we determined that expression of this domain is sufficient to impair the formation of endogenous Dvl1 foci in Nup358-depleted cells (**Fig. 6g, Extended Data Fig. 5d**). These findings indicate that the interaction of Nup358 with Dvl1 through its N-terminal domain prevents the formation of cytoplasmic condensates, inhibiting the degradation of Axin1, and repressing the ligand-independent activation of the Wnt signaling pathway.

## DISCUSSION

Our study establishes Nup358 as a key regulator of intestinal progenitor cells and the development of the intestinal epithelium. We found that by restraining Wnt signaling through the inhibition of Dvl1 condensate formation, Nup358 ensures the generation and survival of transient amplifying progenitors, which give rise to the diverse cell types of the intestinal epithelium and are essential for maintaining crypt-villus architecture. Beyond uncovering a novel role for Nup358 in sustaining intestinal epithelial integrity, our work identified a new molecular mechanism repressing Wnt signaling, which has broad implications for understanding tissue homeostasis and human disease.

Our findings show that Nup358 regulates the function and survival of intestinal progenitor cells. Importantly, in previous work we discovered that in the early stages of hematopoiesis Nup358 is required for the differentiation of myeloid-biased multipotent progenitors and for myeloid development^21^. Moreover, Nup358 was found to play a role in the differentiation of muscle progenitor cells^58^. Altogether, these findings point to Nup358 as a master regulator of stem and progenitor cell biology and a critical determinant of cellular differentiation. Another nucleoporin, Elys, also was linked to intestinal epithelial maintenance in zebrafish and mice^59,60^. ELYS is a member of the essential Nup107-160 NPC scaffold complex and is required for NPC formation. NPC formation is directly linked to cell proliferation, and accordingly, ELYS depletion in zebrafish decreases NPC numbers in highly proliferative tissues, including the intestine. This leads to apoptotic cell death and disrupts tissue integrity^60^. Intestinal-specific ablation of ELYS in mice results in a weaker phenotype compared to zebrafish and, strangely, the intestinal abnormalities resolve in adulthood when knockout mice are almost indistinguishable from controls^59^. While NPC numbers were reported as normal in Elys knockout mice, the data presented does not conclusively rule out defects on NPC assembly. These findings suggest that differently from Nup358, Elys loss affects the survival of proliferating progenitors and the integrity of the intestinal epithelium through its regulation of NPC formation.

Wnt signaling is a central player in intestinal epithelium development and regeneration that controls the balance between stem cell renewal and differentiation^3,30,61,62^. Within the crypt niche, paneth cells and neighboring mesenchymal stromal cells produce Wnt ligands, R-spondins, and other trophic signals that activate canonical Wnt/β-catenin signaling in adjacent intestinal stem cells^3,63,64^. This pathway sustains ISC identity, supports proliferation, and drives the continuous regeneration of the epithelial lining^33,65,66^. Wnt activity is spatially regulated along the crypt-villus axis, forming a gradient that orchestrates epithelial cell fate^33,67^. While high Wnt activity at the crypt base maintains undifferentiated ISCs and fuels TA progenitor expansion, its progressive attenuation toward the villus enforces cell-cycle exit and terminal differentiation into absorptive and secretory lineages^68,69^. Precise modulation of Wnt activity is essential for epithelial turnover both under steady-state conditions and during regeneration, and alterations in this signaling pathway can have profound effects on the integrity of the intestinal epithelium. A moderate increase in Wnt signaling is associated with the expansion of ISCs and TA cells, while excessive or sustained Wnt pathway activation can lead to a differentiation block and trigger apoptosis^35,36^. We found that ablation of Nup358 does not lead to an expansion of the crypt compartment but results in increased levels of activate caspase 3 and the loss of TA cells within the crypts. These findings together with the higher nuclear accumulation of TCF-4 in villus cells indicate that Nup358 loss hyperactivates the Wnt pathway, impairing stem cell differentiation and inducing apoptosis of TA cells and ultimately leading to the depletion of intestinal progenitors.

Previous studies linked Nup358 to canonical Wnt signaling^54,55^. Specifically, the transcription factor TCF-4 was found to interact with several nucleoporins, including Nup358, and to undergo Nup358-dependent SUMOylation^54,55^. TCF-4 SUMOylation was found to enhance its association with β-catenin, favoring its nuclear import^54,55^. These findings exposed a positive role for Nup358 in β-catenin activation. Contrarily, our work identifies Nup358 as a repressor of Wnt signaling, acting upstream of β-catenin/TCF-4 and independently of SUMOylation. We found that loss of Nup358 induces aberrant Dvl1 condensation, triggering Tankyrase-dependent degradation of Axin1. This destabilizes the β-catenin destruction complex, resulting in β-catenin accumulation and ligand-independent activation of Wnt target genes. Echoing our findings, prior work demonstrated an interaction between Nup358 and Dvl1 and aPKC in neurons and implicated Nup358 in restraining Dvl1 activity during neuronal polarity;^42^. The ability of Nup358 to repress cytoplasmic Dvl1 condensate formation is particularly intriguing, given that this nucleoporin harbors multiple phenylalanine-glycine (FG) repeats known to mediate phase separation and has itself been shown to form condensates that regulate nuclear pore complex assembly in early Drosophila embryogenesis^70^. This raises the possibility that Nup358 condensates are physically sequestering Dvl1 away, reducing its free concentration and preventing the nucleation of its own condensates.

Our findings uncover a previously unrecognized mechanism by which cells restrain β-catenin activity, with direct implications for tumorigenesis. Aberrant activation of Wnt/β-catenin signaling is a defining feature of colorectal cancer (CRC), most commonly driven by loss-of-function mutations in core components of the β-catenin destruction complex, APC (∼80-90% of cases) and Axin (5-10%), or stabilizing mutations in β-catenin (∼5-10 %)^71^. While the contribution of Nup358 to cancer remains incompletely understood, accumulating evidence points to a context-dependent role, with both oncogenic and tumor-suppressive functions reported. In CRC patients with a BRAF-like phenotype (∼10–15%), Nup358 supports cell survival by enabling tolerance to mitotic defects in kinetochore-driven microtubule nucleation, consistent with a pro-tumorigenic role^25,72^. In contrast, Nup358 was also shown to suppress tumorigenesis by preserving chromosomal stability through regulation of Topoisomerase IIα SUMOylation^15^. These seemingly opposing roles are likely explained by the distinct functional domains within Nup358, which engage different molecular pathways. Its C-terminal E3 SUMO ligase domain has been implicated in enhancing Wnt signaling by promoting the nuclear accumulation and transcriptional activity of pathway components^54,55^, whereas our work identifies the N-terminal region as a critical repressor of canonical Wnt signaling through inhibition of Dvl1 and promotion of β-catenin degradation. Remarkably, recent genomic analyses of microsatellite instability (MSI) CRC patients identified the N-terminal region of Nup358 as a mutational hotspot, suggesting that disruption of this inhibitory function may be selectively advantageous during tumor evolution^73^. Together, our findings provide a unifying framework to reconcile the dual roles of Nup358 in cancer and position it as a key rheostat of Wnt signaling output. In the context of CRC, where signaling amplitude, rather than binary activation, dictates stemness, differentiation, and tumor progression, loss of Nup358-mediated repression may amplify β-catenin activity beyond a critical threshold, promoting malignant transformation and impairing differentiation programs. These insights not only highlight a new layer of Wnt regulation downstream or parallel to APC loss but also raise the possibility that restoring Nup358 function could represent a targeted therapeutic strategy to dampen oncogenic Wnt signaling in colorectal cancer.

## MATERIALS AND METHODS

### Mice

Nup358 mice were obtained from J. Van Deursen. Animals were bred in a specific pathogen-free facility. All experiments were approved by the Institutional Animal Care and Use Committee of Sanford Burnham Prebys Medical Discovery Institute. To generate Nup358^fl/fl^-CreER^T2^ mice, Nup358^fl/fl^ mice were crossed with R26-CreERT2 mice (The Jackson Laboratory). All mice were used at 9 to 12 weeks of age.

### Antibodies

Antibodies used for immunofluorescence and immunoblotting are as follows: anti-Ki67 (Biolegend, #151206), anti-Nup358/RanBP2 (Bethyl Laboratories, A301-796A), anti-Nup358 (Santa Cruz Biotechnology, sc74518), anti-CD44 (Biolegend, #103012), Anti-activated Caspase 3 (CST #9664). anti-Olfm4 (CST # 39141), anti-TCF-4/TCF7L2 (CST, #2569), anti-Dvl1 (Santa Cruz, #sc8025), anti-Axin1 (CST, #2087), anti-α-Tubulin (CST, #3873), anti-ADP-Ribosylation (CST, #891990), anti-ADP-Ribosylation (CST, #89190), anti-HA (CST, #3724), anti-Histone 3 (CST # 9715), anti-Ubc9 (Abcam, ab33044), anti-Nup98 (Cell Signaling Technology, 2598P), anti-GAPDH (CST, #5174), and anti-GFP (Abcam, #1218).

### Murine primary organoids

Small intestinal organoids were derived from 9–12-week-old *Nup358^fl/fl^-CreER^T^*^2^ and *Nup358^fl/fl^* (control) mice. Mice were euthanized by cervical dislocation, and the proximal small intestine was harvested, flushed with ice-cold phosphate-buffered saline (PBS), and cut longitudinally. Villi were removed by gently scraping the inner side of the intestine with a microscopy slide. The tissue was then cut into approximately 2–5 mm fragments and washed repeatedly in cold PBS until the supernatant ran clear. Intestinal crypts were isolated by four subsequential incubations in cold 5 mM EDTA in PBS at room temperature for 15 minutes with gentle rocking. Each time, after removal of the EDTA solution, small intestinal fragments were subjected to vigorous pipetting in cold 0.1% BSA in PBS. The suspension was filtered through a 70 µm cell strainer to remove villus fragments and debris, and four subsequent enriched crypt fractions were collected by centrifugation at 200 × g for 5 minutes at 4°C. The crypt pellet was washed twice in cold 0.1% BSA in PBS and resuspended in DMEM/F12 and crypt fractions were inspected under the microscope to identify those enriched in crypts and with less villi. After centrifugation, selected crypt pellets were resuspended in warm organoid growth medium (DMEM/F12 supplemented with 10 mM HEPES, 1X GlutaMAX, 1% penicillin/streptomycin, 1X N2 supplement, 1X B27 without vitamin A, 1 mM N-acetylcysteine, 50 ng/mL mouse recombinant EGF, 100 ng/mL Noggin, 1 mM N-acetyl-cysteine, and 10% R-spondin conditioned medium). For plating, equal volumes of crypt suspension and Matrigel were mixed and pipetted into domes in pre-warmed 24-well plates, allowed to polymerize at 37°C for 15–20 minutes, and warm organoid growth medium was then added to each well.

### Cell culture and transfection

293T cells were obtained from American Type Culture Collection (CRL-321) and cultured in humidified conditions at 37°C with 5% CO_2_ in DMEM with 10% heat-inactivated FBS (Sigma-Aldrich). HT-29 cells were obtained from American Type Culture Collection (CRL-321) and cultured in humidified conditions at 37°C with 5% CO_2_ in RPMI with 10% heat-inactivated FBS (Sigma-Aldrich). siRNA transfection was performed with Lipofectamine RNAiMax and cells were analyzed after 6 days. For experiment requiring co-transfection with siRNA and plasmid, cells were first transfected with the siRNA using Lipofectamine RNAiMax, then, after three days, were transfected with the using Lipofectamine 3000, and analyzed 3 days after. siRNAs used in these studies include: control siRNA (Horizon Discovery, D-001810-02-05), Nup358-specific siRNA (Horizon Discovery, J-004746-10-0010), Dvl1-specific siRNA (Horizon Discovery, L-004068-000005), or β-catenin-specific siRNAs (Horizon Discovery, L-003482-00-0005). Plasmids used in this study include β-catenin-EGFP (Addgene, #71367), EGFP-Dvl1 (Addgene, #194586) and plasmids HA-Nup358^1–2683^, HA-Nup358^1–2148^, HA-Nup358^1–1810^, HA-Nup358^1–1133^ from Wälde et al^13^.

### shRNA and viral transduction

Virus was prepared by the Sanford Burnham Prebys Viral Vectors Core. 293T cells were transduced with pTRIPZ tetracycline-inducible short hairpin RNA (shRNA) lentiviruses (control RHS4743 or Nup358 targeting 200761584, 200765221, and 200766435 from Dharmacon), followed by 12 days selection with Puromycin (1.25 μg/ml; Gibco). Expression of shRNA was induced with doxycycline (200 ng/ml; Clontech). To generate a stable Wnt reporter line, HEK293T cells were transduced with lentivirus derived from the 7TGP plasmid encoding GFP under the control of seven tandem TCF/LEF binding sites (Addgene #24305). Following transduction, cells were maintained under puromycin selection (1.25 μg/mL; Gibco) for 12 days. Reporter activity was validated by treating the stable line with recombinant Wnt3a (100 ng/mL, Proteintech HZ-1296) and confirming EGFP induction via fluorescence microscopy.

### XAV-939, 2-D08, and 1,6-hexanediol treatments

For pharmacological inhibition of Tankyrase activity, the culture media was replaced with media containing 200nM XAV-939 or DMSO vehicle control at 72 hours post-siRNA transfection, and cells were incubated for additional 72 hours prior to analysis. For SUMOylation inhibition, cells were treated with 25 μM 2-D08 (Selleckchem, S8696) 72 hours after transfection with either control or Nup358 siRNA. Treatment was maintained for 24 hours prior to analysis as previously described^21^. Vehicle controls consisted of equivalent volumes of DMSO. To disrupt biomolecular condensates, cells transfected with Dvl1-EGFP were treated with 5% 1,6-hexanediol (v/v in complete medium) for 2 minutes at 37°C. Cells were fixed for 10 minutes with 4% PFA and analyzed by fluorescent microscopy.

### Immunofluorescence Staining

Cells were fixed in 4% paraformaldehyde (PFA) for 5 minutes at room temperature (RT). Pre-hybridization was performed in immunofluorescence (IF) buffer [PBS, 1% bovine serum albumin (BSA), 0.02% SDS, and 0.1% Triton X-100] for 30 minutes at RT. Primary antibodies were incubated overnight at 4°C. Cells were washed three times with IF buffer, incubated with secondary antibodies, labeled with Hoechst 33342, and washed with PBS. For organoid immunofluorescence staining, organoids were fixed in 4% PFA in PBS for 30 minutes at room temperature, followed by three 5 minutes washes in PBS. Fixed organoids were permeabilized with 0.5% Triton X-100 in PBS for 10 minutes at RT, and three times in 100mM glycine in PBS for 10 minutes. Non-specific binding was blocked by incubating organoids in blocking buffer made of 10% goat serum diluted in IF buffer for 1 hour at RT. Primary antibodies were diluted in blocking buffer and incubated with the organoids overnight at 4°C. Organoids were washed three times in IF buffer for 20 minutes and incubated with secondary antibodies and Hoechst in blocking buffer for 1 hour at RT. Organoids were washed three times for 20 minutes in IF buffer and mounted using ProLong Glass Antifade Mountant (Invitrogen 2342881). Fluorescence imaging was performed with DMi8 Leica SP8 confocal microscope and analyzed in Leica Application Suite X software v3.1.5.16308.

### Live imaging and fluorescence recovery after photobleaching (FRAP)

For live imaging, cells and organoids were seeded in polymer coverslip 8-well chamber slides (Ibidi). At the desired time of analysis, cells were transferred in a humidified 37°C with 5% CO_2_ stage top incubator (H301, Okolab) controlled by the Oko-Touch (Okolab) and imaged on a Leica DMi8 confocal microscope. For FRAP, 293t cells stably expressing Wnt reporter were seeded in polymer coverslip 8-well chamber slides and transfected with control or Nup358 siRNAs. After 6 days from transfection, cells were imaged on a Leica DMi8 confocal microscope. EGFP-positive signal from cytoplasmic foci was photobleached with 50% laser power for 3 seconds. Recovery was recorded for 20 seconds using the LAS X software.

### Flow cytometry

For flow cytometric quantification of 7TGP Wnt reporter activation, cells were harvested, stained with Zombie NIR Fixable Viability Kit to allow for live/dead discrimination. Analytical cytometry was performed in the Sanford Burnham Prebys Flow Cytometry Core using a BD LSRFortessaTM (BD Biosciences) or a BD LSRFortessaTM X-20 (BD Biosciences). GFP fluorescence was measured with gates established using untransfected and single-fluorochrome controls.

### qPCR

Small intestinal epithelium was harvested from *Nup358^fl/fl^-CreER^T^*^2^ and *Nup358^fl/fl^* (control) mice and RNA was isolated with Trizol according to manufacturer instructions. cDNA was synthesized using the QuantiTect Reverse Transcription Kit (QIAGEN). qPCR was carried out using SYBR Green (ThermoFisher Scientific). qPCR data were collected in a CFX384 Real-Time PCR Detection System (Bio-Rad). The following primers were used to detect mouse transcripts: Axin2 5’-GGCCCTGCTGTAAAAGAGAGGA-3’ (forward) and 5’-TTCCTCTCAGCAATCGGCGT-3’ (reverse); Ccnd1 5’-TCAAGTGTGACCCGGACTGC-3’ (forward) and 5’-TGGGGTCCATGTTCTGCTGG-3’ (reverse); cMyc: 5’-GTGGAAAACCAGCAGCCTCC-3’ (forward) and 5’-GCACCGAGTCGTAGTCGAGG-3’ (reverse). Expression was normalized to ACTB as housekeeping genes and analyzed by the 2−ΔΔCt method.

### Western blotting

For protein extracts, cells were washed with ice-cold PBS containing Protease and Phosphatase inhibitors (Pierce Halt Protease and Phosphatase Inhibitor Cocktail, ThermoFisher Scientific) and harvested with RIPA buffer containing Protease and Phosphatase inhibitors. Homogenates were incubated on ice for 30 minutes and passed 5-10 times though a 29 G syringe to shear the DNA. Protein concentration was determined using the Pierce BCA reagent (ThermoFisher Scientific). LDS Sample Buffer premixed with NuPAGE® Sample Reducing Agent (Life Technologies) was added, and samples were incubated for 10 minutes at 70°C. For western blot analysis, 20 or 40 µg of protein were resolved by SDS-PAGE on NuPAGE Novex 3-8% tris-acetate protein gels or Bolt 4-12% Bis-Tris Plus protein gels (Invitrogen) and blotted to nitrocellulose membranes using an iBlot2 Dry Blotting System. Membranes were stained with Ponceau, washed with TBS, and blocked for 1 hour at RT with TBS-0.1% Tween 20 (TBS-T) containing 5% BSA (for phospho-antibodies) or 5% nonfat milk, and incubated with primary antibodies overnight at 4°C. After three washes with TBS-T, the secondary antibody was added and incubated for 1 hour at RT. Membranes were then washed and developed using the Pierce ECL Western Blotting Substrate (ThermoFisher Scientific) or SuperSignal West Pico Plus Chemiluminescent Substrate (ThermoFisher Scientific).

### Co-immunoprecipitation

To investigate the interaction between Nup358 and Dvl1, co-immunoprecipitation studies were performed. 293T cells were harvested and washed twice with PBS and homogenized in RIPA buffer containing HALT Protease and Phosphatase Inhibitor Cocktail (Thermo Fisher Scientific). DNA was sheared using a 29G syringe. Samples were centrifuged at 3000g at 4°C for 10 minutes to remove debris. Protein concentration was measured using the Pierce bicinchoninic acid (BCA) reagent. Eight milligrams of total proteins were diluted to the final concentration of 2 mg/ml in immunoprecipitation (IP) buffer [50 mM Tris (pH 7.4), 150 mM NaCl, 0.5% NP-40, 1 mM EDTA, 1 mM MgCl2, and 1 mM dithiothreitol] containing protease and isopeptidase inhibitors and incubated with Protein A/G Magnetic Beads (Thermo Fisher Scientific) for 2 hours to remove nonspecific binding. Precleared protein solution was incubated over-night at 4°C on a rotator with either a specific anti-Dvl1 antibody (Santa Cruz #8025) at the final dilution of 1:400 or with the same amount of control rabbit IgG. The next morning, Protein A/G Magnetic Beads were added, and samples were incubated on a rotator for 4 hours at 4°C. After collection of the flow through, immunocomplexes were washed three times with IP buffer and eluted from the magnetic beads by incubation with NuPAGE LDS sample buffer (Thermo Fisher Scientific) with reducing agent for 15 minutes at 70°C and analyzed by immunoblot with an anti-Nup358 antibody (Santa Cruz Biotechnology, sc74518). Similar co-immunoprecipitation studies were performed in 293T cells harvested after 72 hours from transfection with Dvl1-EGFP and HA-Nup358^1–1133^. Immunoprecipitation was performed with an anti-GFP antibody (Abcam, #1218) and western blot was carried out with an antibody against HA tag (CST, #3724).

### Data collection and analysis

GraphPad Prism 8 software v8.2 (GraphPad Software, Inc.) was used to prepare graphs and to perform statistical analyses. Student’s t test was applied when comparing two groups. Multiple t tests with corrections for multiple comparisons or two-way ANOVA were used when comparing greater than two groups. Microscopy data were collected with a DMi8 Leica SP8 confocal microscope and analyzed using the Leica Application Suite X software v3.1.5.16308 (Leica Microsystems) or ImageJ v2.0.0-rc-54/1.51h (NIH) or Fiji (NIH). Flow cytometry data were collected using the BD FACSDIVATM Software (BD Biosciences) and analyzed using FlowJo software v10.0.8r1 (Tree Star, Inc.). qPCR data were collected in a CFX384 Real-Time PCR Detection System (Bio-Rad).

## Acknowledgements

We thank Dr. Jan Van Deursen for kindly providing the Nup358^fl/fl^ mouse line, Dr. Ralph Kehlenbach for providing HA-Nup358 expression plasmids, and Dr. Michael Cohen for providing the anti-ADP-ribosylation antibody. This work was supported by the NCI Cancer Center grant P30 CA030199, which supports the animal, flow cytometry, genomics and bioinformatics cores at the SBP La Jolla campus.

## Author contributions

V.G. designed experimental approach, performed experiments, analyzed data and co-wrote the manuscript; S.S. designed experimental approach, performed experiments, and analyzed data; E.Y.S.Z. assisted with experiments; D.L. assisted with experiments; M.A.D. designed experimental approach, analyzed data, provided oversight and critical expertise, and co-wrote the manuscript.

