## Supplemental Figures 1-5 for "Nup358 Sustains Intestinal Epithelial Homeostasis by Preventing Dvl1 Condensate Formation to Restrain Wnt Signaling" (from another version of this preprint: version 1, 2026-03-26)

### Extended Data Figure 1

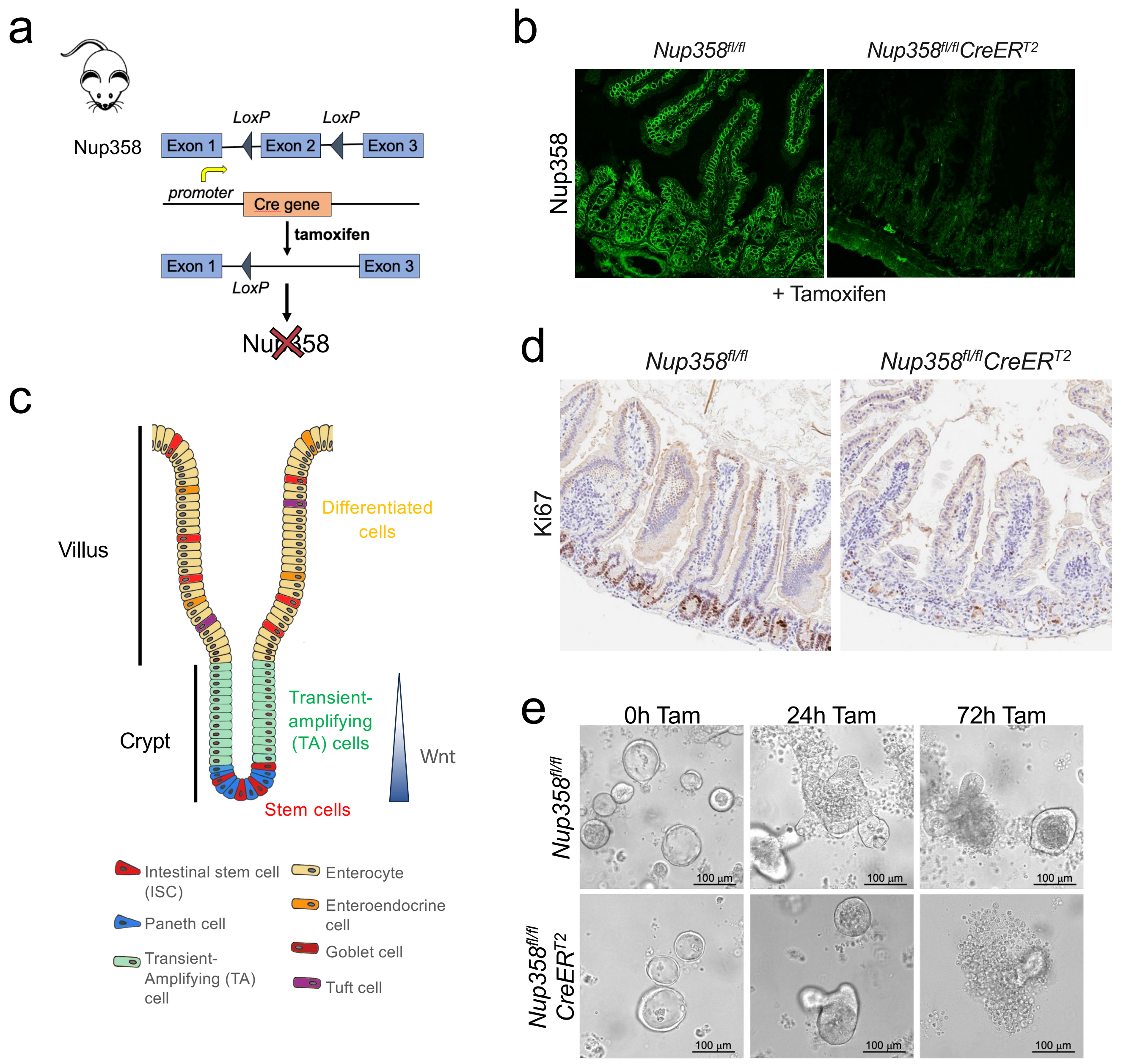

**Extended Data Figure 1. Nup358 ablation in the intestinal epithelium and organoids.**

**a.** Schematic illustration of Nup358 knockout strategy.

**b.** Immunofluorescence staining for Nup358 in small intestinal epithelium of *Nup358*$^{fl/fl}$ and *Nup358*$^{fl/fl}$ *CreER*$^{T2}$ mice treated with tamoxifen.

**c.** Schematic illustration of the small intestinal epithelium architecture and domains.

**d**. Immunohistochemical staining for Ki67 in small intestinal epithelium from *Nup358*$^{fl/fl}$ and *Nup358*$^{fl/fl}$ *CreER*$^{T2}$ mice treated with tamoxifen.

**e**. Intestinal organoids (IOs) generated from crypts isolated from *Nup358*$^{fl/fl}$ and *Nup358*$^{fl/fl}$ *CreER*$^{T2}$ mice were cultured *in vitro*. Tamoxifen was added after 24 hours of culture. Images show organoids before adding tamoxifen (0h) and at 24h and 72h after tamoxifen addition.

### Extended Data Figure 2

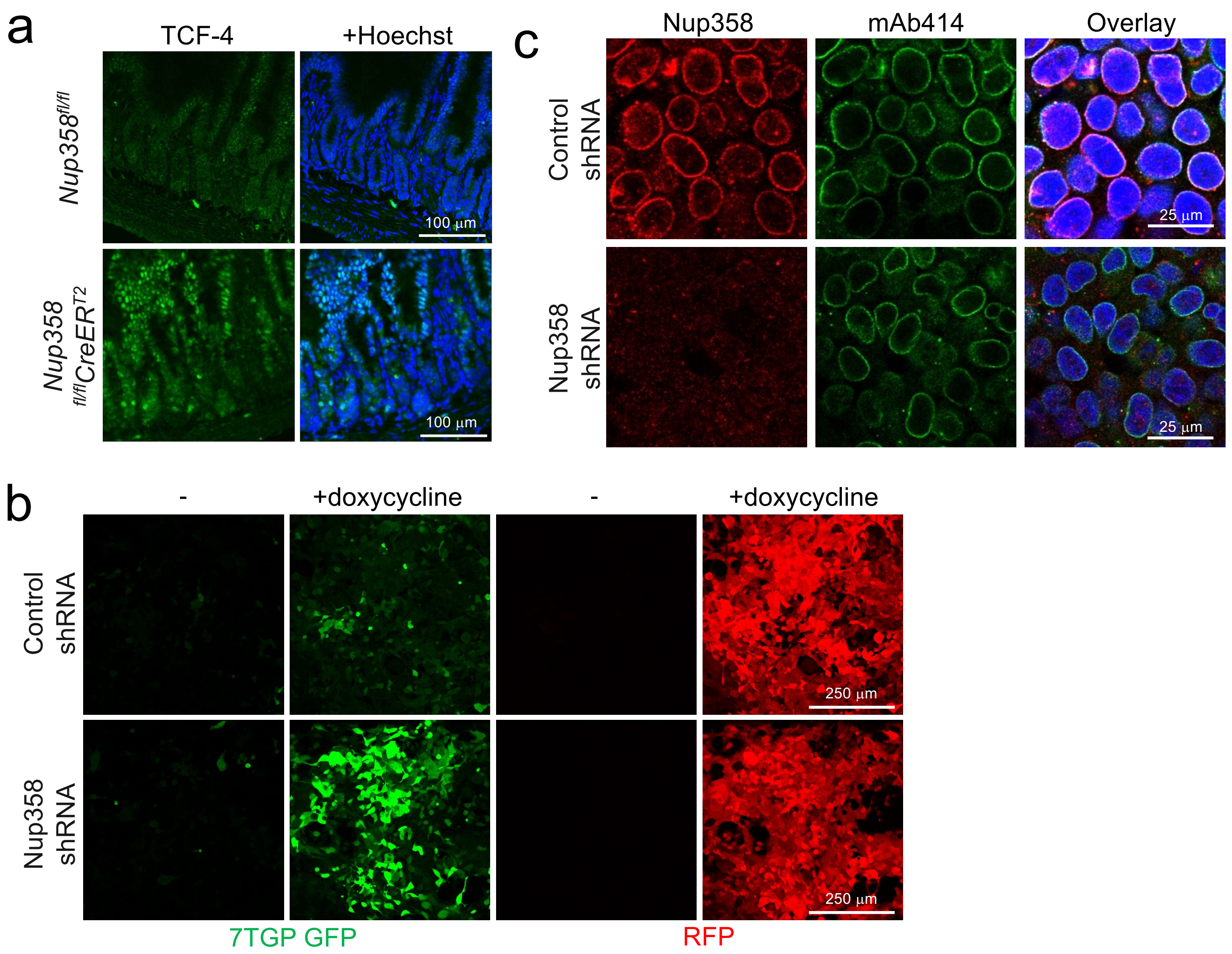

**Extended Data Figure 2. Wnt signaling activation by Nup358 depletion.**

**a.** Immunofluorescence staining for TCF-4 in small intestinal epithelium from *Nup358*$^{fl/fl}$ and *Nup358*$^{fl/fl}$ *CreER*$^{T2}$ mice treated with tamoxifen.

**b**. Immunofluorescence staining for Nup358 and the NPC marker mAb414 of HEK293T cells expressing control or Nup358-specific shRNAs.

**c**. Confocal microscopy analysis of 7TGP Wnt reporter expression levels (GFP) and shRNA expression levels (RFP) of HEK293T cells expressing control or Nup358-specific shRNAs.

### Extended Data Figure 3

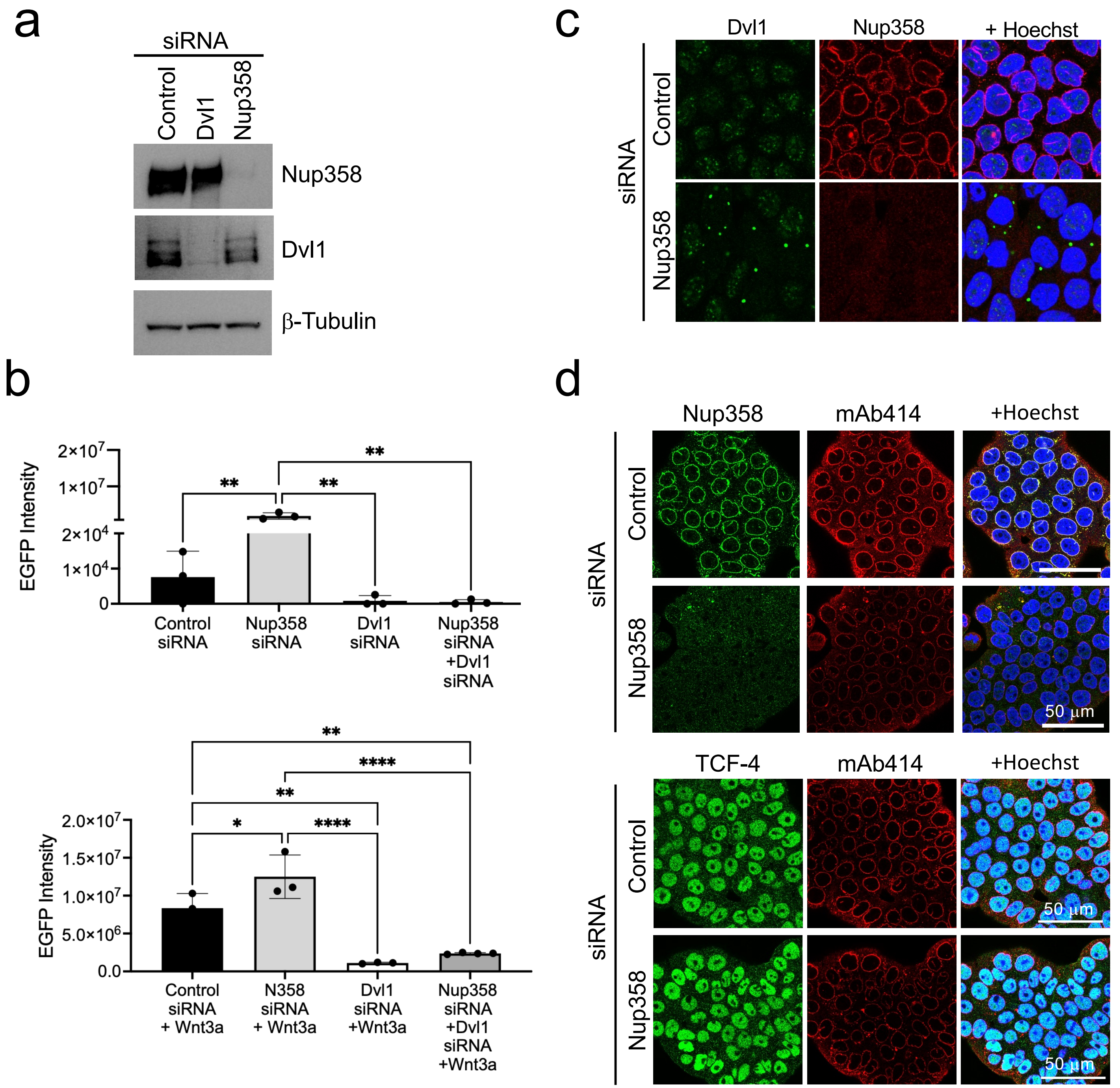

**Extended Data Figure 3. Nup358 functions through Dvl1**

**a.** Western blot analysis of Nup358 and Dvl1 in HEK293T cells transfected with control, Dvl1-, and Nup358-specific siRNAs. β-Tubulin was used as loading control.

**b.** Quantification of 7TGP fluorescent Wnt reporter activity in HEK293T cells transfected with control, Dvl1-, or Nup358-specific siRNAs, or a combination of Nup358- and Dvl1-specific siRNAs and treated with either Wnt3a or vehicle. Data are expressed as mean ± SD., $^{*}p \leq 0.05$, $^{**}p \leq 0.01$, $^{****}p \leq 0.0001$.

**c**. Immunofluorescence staining for Dvl1 and Nup358 in HT-29 colorectal cancer cells transfected with control or Nup358-specific siRNAs.

**d.** Immunofluorescence staining for Nup358 and TCF-4 in HT-29 colorectal cancer cells transfected with control or Nup358-specific siRNAs. mAb414 was used to mark nuclear pore complexes.

### Extended Data Figure 4

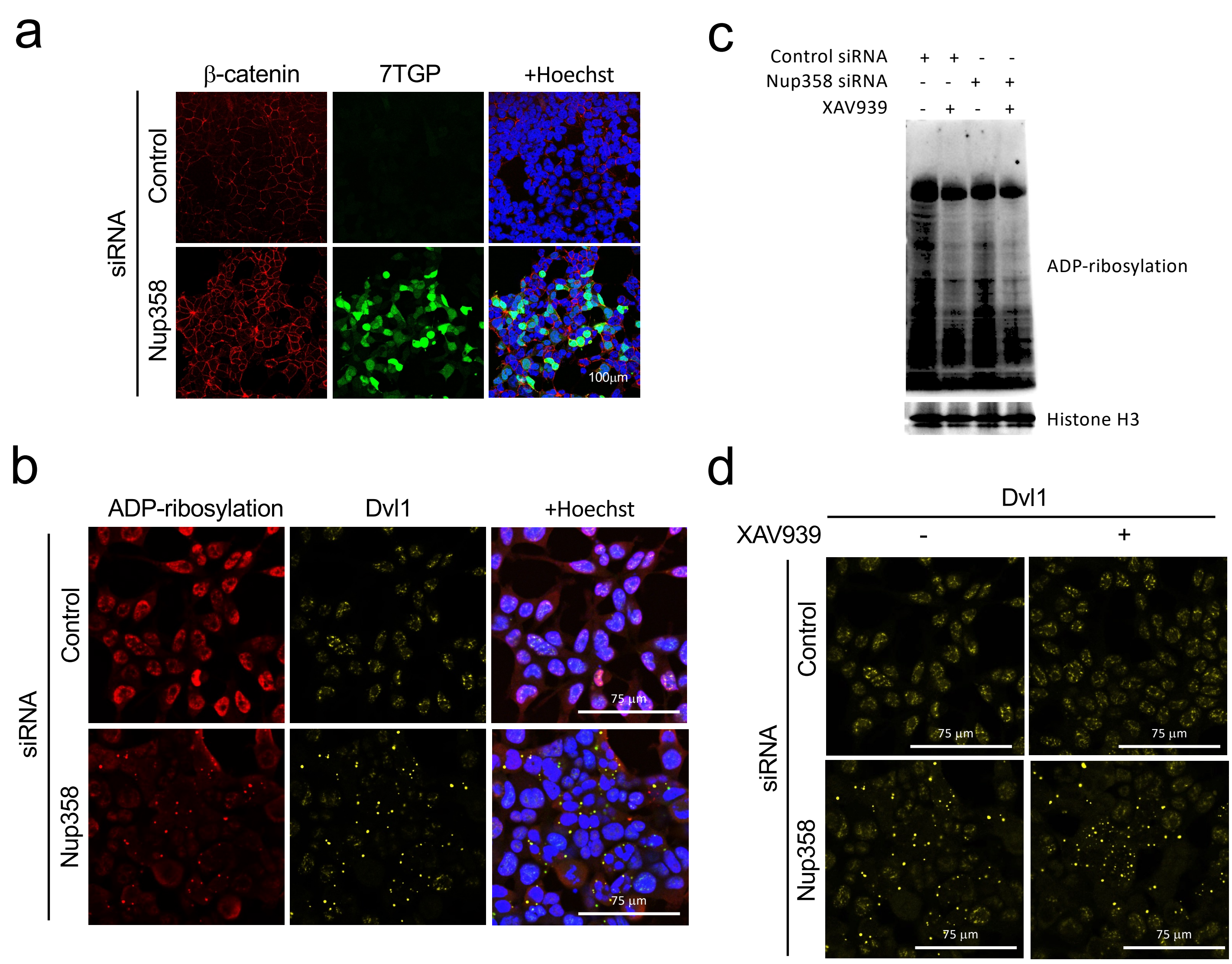

**Extended Data Figure 4. Nup358 depletion stabilizes β-catenin and affects ADP-ribosylation.**

**a.** Immunofluorescence staining for β-catenin in HEK293T cells stably expressing 7TGP Wnt reporter and transfected with control or Nup358-specific siRNAs.

**b.** Immunofluorescence staining for ADP-ribosylation and Dvl1 in HEK293T cells transfected with control or Nup358-specific siRNAs.

**c**. Western blot analysis of ADP-ribosylation in HEK293T cells transfected with control siRNA or Nup358-specific siRNAs in the presence or absence of the ADP-ribosylation inhibitor XAV939. Histone 3 was used as loading control.

**d**. Immunofluorescence staining for Dvl1 in HEK293T cells transfected with control or Nup358-specific siRNAs and treated with either vehicle or ADP-ribosylation inhibitor XAV939.

### Extended Data Figure 5

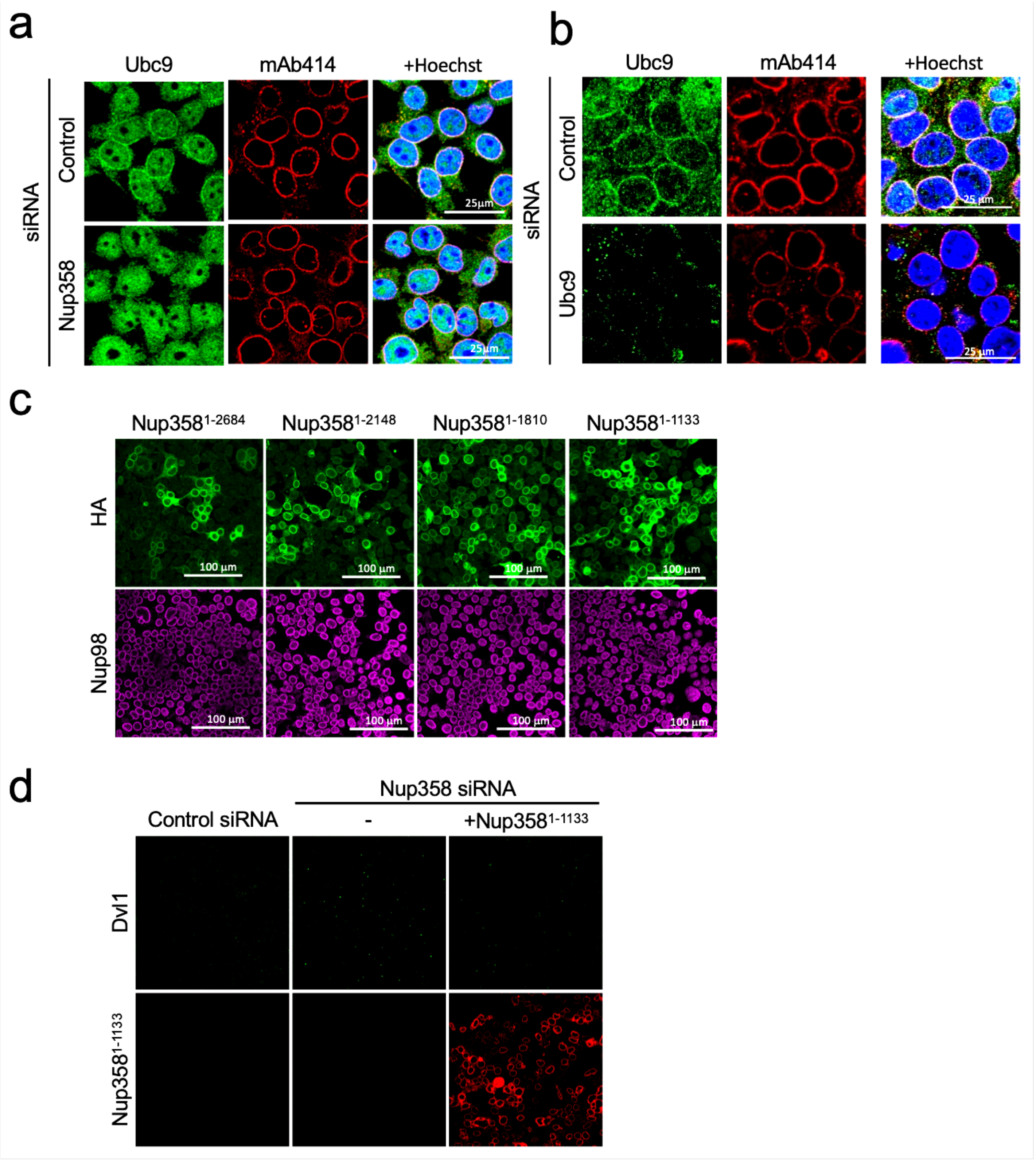

**Extended Data Figure 5. Nup358 depletion impairs Ubc9 NPC localization and Nup358 N-terminal inhibits Dvl1 foci formation.**

**a.** Immunofluorescence staining for Ubc9 in HEK293T cells transfected with control or Nup358-specific siRNAs. mAb414 was used to mark nuclear pore complexes.

**b.** Immunofluorescence staining for Ubc9 in HEK293T cells transfected with control or Ubc9-specific siRNAs. mAb414 was used to mark nuclear pore complexes.

**c.** Immunofluorescence staining of HEK293T cells expressing different HA-tagged truncated forms of Nup358. Nup98 was used to mark nuclear pore complexes.

**d.** Immunofluorescence staining for Dvl1 and HA in HEK293T cells transfected with control siRNA, Nup358 siRNA, or a combination of Nup358 siRNA and HA-tagged $Nup358^{1-1133}$.

**Extended Data Video 1.** Live imaging of intestinal organoids generated from $Nup358^{fl/fl}$ (control) mice after 96h of tamoxifen treatment. Imaging was performed for 4 days.

**Extended Data Video 2**. Live imaging of intestinal organoids generated from $Nup358^{fl/fl}$ $CreER^{T2}$ (Nup358 KO) mice after 96h of tamoxifen treatment. Imaging was performed for 4 days.

**Extended Data Video 3.** Fluorescence recovery after photobleaching (FRAP) analysis in HEK293T cells transfected with Nup358-specific siRNA and EGFP-Dvl1.
